# Detection of merbecoviruses in Danish bats using a novel broad-range RT-qPCR assay

**DOI:** 10.64898/2026.10.20.760357

**Authors:** Camille Melissa Johnston, Ana Moreno, Davide Lelli, Elinor Lindberg Marving, Hans J. Baagøe, Morten Elmeros, Esben T. Fjederholt, Louise Lohse, Thomas Bruun Rasmussen

## Abstract

Bat-associated merbecoviruses comprise a genetically diverse group of coronaviruses with variable receptor usage and host range, highlighting the need for efficient tools for their surveillance. Here, we developed and optimized a broad-range reverse transcription quantitative PCR (RT-qPCR) assay for the detection of bat-associated merbecoviruses, including the lineage previously identified in brown long-eared bats (*Plecotus auritus*) in Denmark. *In silico* analysis predicted detection of 898 of 941 (95.4%) merbecovirus sequences, and the assay detected all bat-associated merbecoviruses included in the experimental panel. The assay was applied to 33 bats sampled through the Danish national bat surveillance program in 2025. Merbecovirus RNA was detected in two samples, one from a *P. auritus* and one from a Nathusius’ pipistrelle (*Pipistrellus nathusii*), representing the first detection of a merbecovirus in *P. nathusii* in Denmark. Metagenomic next-generation sequencing recovered a complete merbecovirus genome from the *P. nathusii* sample. Phylogenetic analysis assigned the virus to the MOW clade and showed that it was closely related to the Russian MOW15-22 virus, sharing 98.67% nucleotide identity across the complete genome and 99.85% amino acid identity in the spike protein. Limited sequence data obtained from the *P. auritus* sample was consistent with the merbecovirus lineage previously identified in this species in Denmark. Together, these findings demonstrate the utility of broad-range RT-qPCR combined with metagenomic sequencing for surveillance and genomic characterization of bat-associated merbecoviruses.

## 1. Introduction

Coronaviruses (CoVs) are enveloped RNA viruses associated with a wide range of respiratory and enteric diseases in humans and animals (Corman et al., 2018). Bats (order Chiroptera) are important reservoirs of mammalian coronaviruses, harboring the greatest diversity of CoVs among all known host groups (Hu et al., 2015; Letko et al., 2020). They are also regarded as the evolutionary source of several zoonotic coronaviruses, including severe acute respiratory syndrome (SARS-CoV) and SARS-CoV-2 (Hu et al., 2015; Temmam et al., 2022), and possibly Middle East respiratory syndrome coronavirus (MERS-CoV) (Anthony et al., 2017; Memish et al., 2013).

Coronaviruses have positive-sense RNA genomes of approximately 27-31 kb (de Groot et al., 2012). The 5’ end of the genome encodes the replicase complex, comprising the overlapping open reading frames ORF1a and ORF1b, which are translated through a ribosomal −1 frameshift into polyproteins that are subsequently cleaved into 16 non-structural proteins (Sola et al., 2015). The 3’ region of the genome encodes the structural proteins, including the spike (S) protein, and a variable number of accessory genes, which differ among coronavirus lineages (van Boheemen et al., 2012).

The genus *Betacoronavirus* comprises several subgenera of importance to human and animal health, including *Sarbecovirus*, which contains SARS-CoV and SARS-CoV-2, and *Merbecovirus*, containing MERS-CoV (Woo et al., 2023). The latter has attracted particular attention following the emergence of MERS-CoV and the growing recognition of a diverse reservoir of related viruses in wildlife. Merbecoviruses have been identified in a wide range of mammalian hosts across Africa, Europe, Asia and the Americas, with bats representing their primary reservoir (Chen et al., 2023; De Sabato et al., 2020; Haagmans et al., 2014; He et al., 2022; Ithete et al., 2013; Lau et al., 2013; Luo et al., 2018; Moreno et al., 2017; Wang et al., 2014; Y. Yang et al., 2014). In particular, numerous merbecoviruses have been detected in insectivorous bats of the family *Vespertilionidae*, including species belonging to the widespread genera *Myotis*, *Pipistrellus*, *Plecotus*, and *Eptesicus* (Annan et al., 2013; Anthony et al., 2017, 2013; Corman et al., 2014; Ithete et al., 2013; Lelli et al., 2013; L. Yang et al., 2014). Despite the increasing number of merbecoviruses identified in vespertilionid bats, their diversity, geographic distribution, and evolutionary relationships remain incompletely characterized in many regions.

Recent phylogenetic analyses based on receptor-binding domains (RBDs) have revealed substantial diversity within the subgenus *Merbecovirus*, identifying at least eight major clades: EriCoV, NeoCoV, MOW, HKU5-1, HKU5-2, HKU25, HKU4 and MERS (Catanzaro et al., 2025; Chen et al., 2025; Park et al., 2025). This diversity is accompanied by marked variation in receptor usage. Members of the MERS and HKU4 clades utilize dipeptidyl peptidase IV (DPP4) for cell entry, whereas viruses belonging to the NeoCoV, MOW, HKU5, and HKU25 clades have been shown to utilize angiotensin-converting enzyme 2 (ACE2), albeit with differing host specificities (Catanzaro et al., 2025; Chen et al., 2025; Liu et al., 2025; Ma et al., 2025, 2023; Park et al., 2025; Xiong et al., 2022). More recently, aminopeptidase N (APN) was identified as the entry receptor for members of the EriCoV clade (Jin et al., 2025). Together, these findings highlight substantial evolutionary and functional diversity among bat-associated merbecoviruses.

We recently reported the first detection of merbecoviruses in Denmark, recovering two complete genomes from brown long-eared bats (*Plecotus auritus*) that were assigned to the HKU25 clade by phylogenetic analysis (Johnston et al., 2025). However, the broader diversity and distribution of merbecoviruses among Danish bat populations remain largely unknown. Improved surveillance tools are therefore needed to determine the occurrence and genetic diversity of these viruses across bat species.

Metagenomic next-generation sequencing (mNGS) have greatly enhanced virus discovery by enabling untargeted detection and genomic characterization of viruses present in complex biological samples (Nooij et al., 2018). However, the cost and analytical requirements can limit its application for large-scale routine surveillance programs. Targeted RT-qPCR assays provide a rapid, sensitive, and cost-effective approach for screening large sample collections, with positive samples subsequently subjected to mNGS for more detailed genomic characterization. Combining these approaches therefore provides an efficient strategy for targeted surveillance and characterization of bat-associated coronaviruses.

In this study, we developed and optimized a broad-range RT-qPCR assay targeting for the detection of bat-associated merbecoviruses, including the *P. auritus*-associated lineage identified previously in Denmark. We applied the assay to samples collected through the Danish bat surveillance program and characterized the RT-qPCR positive samples by mNGS. This approach identified merbecoviruses in two bat species. A virus detected in *P. auritus* was closely related to the merbecovirus previously reported from this host in Denmark, whereas a merbecovirus detected in Nathusius’ pipistrelle (*Pipistrellus nathusii*) represented a distinct merbecovirus lineage not previously detected in Denmark. The resulting genomes were analyzed to determine their phylogenetic relationships and evolutionary placement within the subgenus *Merbecovirus*.

## 2. Methods

### 2.1. Primer design and assay development

Sequences belonging to the previously described bat-associated merbecovirus clade (Johnston et al., 2025), together with the closely related MOW15-22 sequence (GenBank accession ON325306.1), were aligned using MAFFT v7.526 (Katoh and Standley, 2013) with default parameters (**Table S1**). Primers targeting conserved regions within ORF1ab were initially designed manually using only sequences from the bat-associated merbecovirus clade (**Table 1**). Additional primers were generated using our semi-automated primer design workflow (Johnston et al., 2026b), which iteratively runs varVAMP (Fuchs et al., 2025) across user-defined consensus thresholds and maximum primer degeneracy levels. varVAMP designs degenerate primers for (RT)-qPCR based on a multiple sequence alignment. For this design, the complete alignment, including MOW15-22, was used as input. Consensus thresholds of 0.90 and 0.95 and maximum primer degeneracy levels of 3–6 ambiguous positions were evaluated to generate candidate assays with broad sequence coverage (**Table 1**).

**Table 1:**
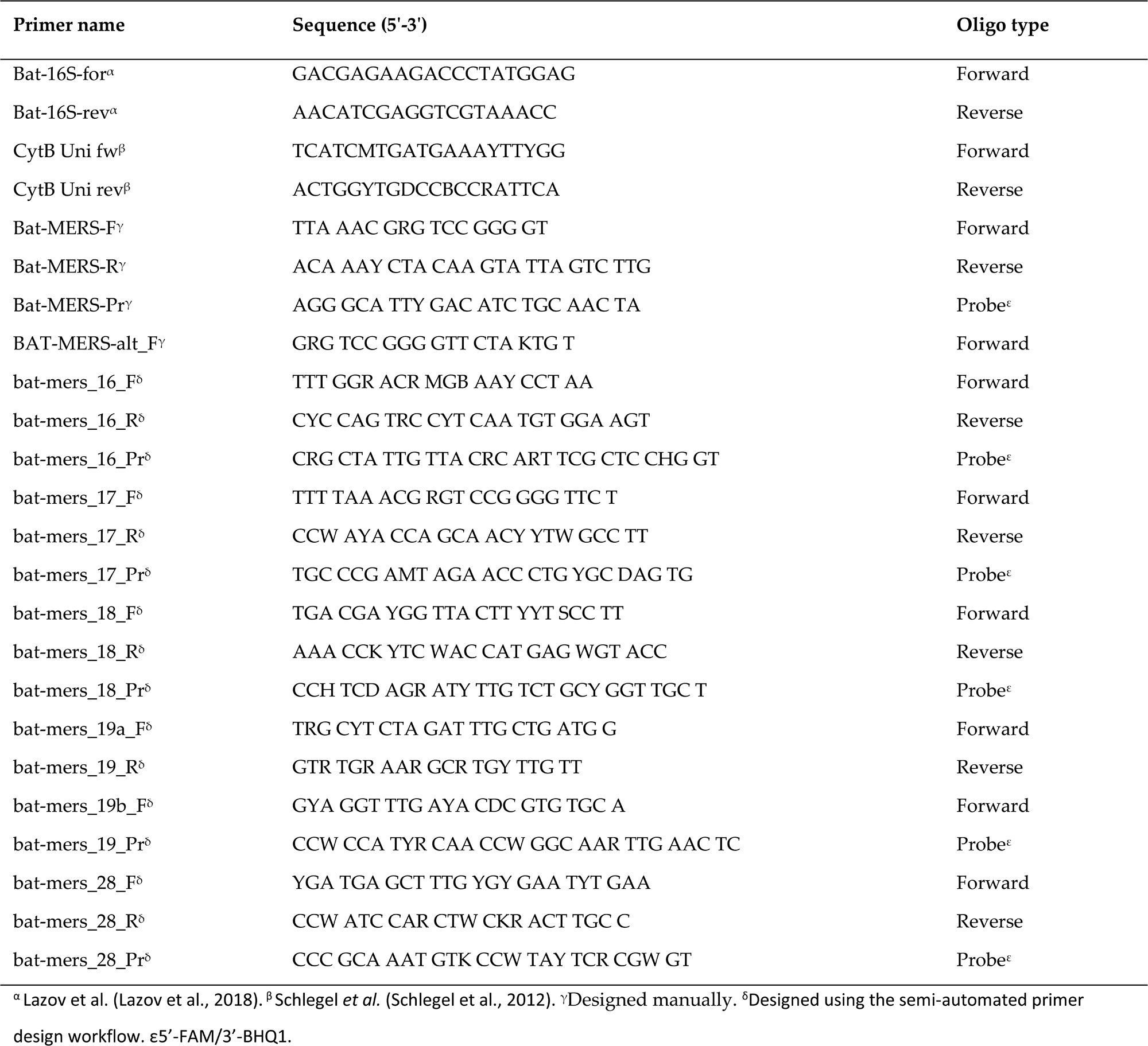
Primer overview.

Primer and probe sets were initially evaluated using purified RNA from a 100-fold dilution of a merbecovirus-positive sample (BtCoV/228741-35/H.sav/IT/2019). RT-qPCR reactions were performed using the Qiagen OneStep RT-PCR kit (QIAGEN, Hilden, Germany) in a final reaction volume of 12.5 µl, with primer and probe concentrations of 0.4 µM and 0.2 µM, respectively. Thermal cycling conditions consisted of using 50°C for 30 min, followed by 95°C for 15 min, then 45 cycles of 95°C for 15s, 55°C for 30s and 72°C for 30s, with a final step at 25°C for 2 min.

Selected assays were subsequently used to compare the performance of three one-step RT-qPCR kits: Qiagen OneStep RT-PCR kit (Qiagen), AgPath ID One-Step RT-PCR (ThermoFisher Scientific, Waltham, MA, USA) and SuperScript III One-Step RT-PCR (ThermoFisher Scientific). For the Qiagen kit, thermal cycling was performed as above. For AgPath ID, cycling conditions consisted of 50°C for 10 min; 95°C for 10 min; 45 cycles of 95°C for 15 s, 55°C for 20 s, 72°C for 30 s. For SuperScript III, cycling conditions consisted of 50 °C for 30 min; 94°C for 2 min; 45 cycles of 94°C for 15 s, 55°C for 30 s, 68°C for 30 s. Both protocols included a final step at 25°C for 2 min.

Finally, selected assays were validated against a panel of bat- and hedgehog-associated merbecoviruses as well as non-target coronaviruses, including bat alphacoronaviruses and a bat sarbecovirus (**Table S2**). The panel was used to assess analytical reactivity within the target group and potential cross-reactivity with non-target coronaviruses.

### 2.2. *In silico* primer analysis

Selected assays were evaluated using our previously described primer prediction workflow (Johnston et al., 2026b), which uses BLAST (Camacho et al., 2009) to predict on- and off-target primer capture against local sequence databases. Two databases were generated from the previously described *Coronaviridae* dataset (Johnston et al., 2026a): One containing all *Coronaviridae* sequences excluding *Merbecovirus*, and one containing only *Merbecovirus* sequences. The *Merbecovirus* database was used to assess predicted assay inclusivity, whereas the non-Merbecovirus *Coronaviridae* database was used to evaluate potential cross-reactivity.

### 2.3. Collection of samples

Fecal samples from live bats and fecal or colon samples from bat carcasses representing nine species were collected as part of a national surveillance program of bats in Denmark in 2025 (**Table S5**). The program included both active surveillance, based on non-invasive collection of fecal samples from live bats, and passive surveillance, based on sampling of carcasses submitted from multiple geographic regions across Denmark. Sampling from live bats was conducted with minimal disturbance to the animals. Metadata, including species identification, sampling location, and date of collection, were systematically recorded for each sample.

### 2.4. Host species identification

Live bats were identified to species level based on morphological characteristics. Bats obtained through passive surveillance were identified by PCR amplification and sequencing of mitochondrial 16S rRNA and cytochrome b genes (**Table 1**), as previously described (Lazov et al., 2018).

### 2.5. Screening of samples using a novel merbecovirus RT-qPCR assay

Fecal and colon samples were homogenized in Eagle’s medium using a TissueLyser II (QIAGEN) and clarified by centrifugation. The RNA was extracted from the resulting supernatant using a MagNA Pure 96 system (Roche, Basel, Switzerland) with the DNA and Viral NA Small Volume Kit and the Viral NA plasma external lysis S.V. 3.1 protocol. Extracted RNA was screened for merbecoviruses using the bat-MERS-F_17Pr-R RT-qPCR assay developed in this study. Reactions were performed using the Qiagen OneStep RT-PCR kit under the conditions described above.

### 2.6. Genome sequencing and assembly

Samples positive in the merbecovirus RT-qPCR assay were subjected to metagenomic sequencing. Double-stranded cDNA was generated as previously described (Lazov et al., 2021) and sequencing libraries were prepared as previously described (Johnston et al., 2025). Libraries were sequenced on an Illumina MiSeq platform (Illumina, San Diego, CA, USA), generating 2×150 bp paired-end reads.

Sequence reads were processed using a previously described metagenomics workflow (Johnston et al., 2026a). Briefly, reads were quality-trimmed, and viral and unclassified reads were extracted for *de novo* assembly. Draft genomes were generated and subsequently refined by manual inspection. Genome annotation was performed by reference-based transfer followed by manually curation, including identification of the ORF1a/ORF1ab ribosomal frameshift site.

For samples with insufficient reads for *de novo* genome assembly, reference-based mapping was performed as previously described (Johnston et al., 2026a). A partial consensus sequence was subsequently generated in Geneious Prime 2026.0.1 (Biomatters Inc., Boston, MA, USA), retaining positions with a coverage depth ≥1, while positions without read coverage were represented as gaps.

### 2.6. Phylogenetic analysis

To place the complete *P. nathusii*-associated genome within the broader diversity of coronaviruses, alignment-free pairwise distances based on total average nucleotide identity (tANI) were computed using Vclust v1.3.1 (Zielezinski et al., 2025) against a previously described representative CoV dataset (Johnston et al., 2026a) (**Table S6**), including novel sequences. A neighbor-joining tree was subsequently constructed from the pairwise distance matrix as previously described (Johnston et al., 2026a).

To resolve phylogenetic relationships within the subgenus *Merbecovirus*, beta-CoV sequences were extracted from the CoV dataset described in Johnston et al. 2026, and supplemented with recently available genomes (excluding *Betacoronavirus pandemic*) from NCBI (accessed 2026-05-13). A representative dataset was generated using Vclust at 81% tANI with subclustering at 91% tANI, limiting clusters to approximately 10 genomes, as described previously (Johnston et al., 2026a). Merbecovirus sequences were subsequently extracted to generate the final dataset (**Table S7**).

Sequences generated in this study were aligned with the representative dataset using MAFFT v7.526 (Katoh and Standley, 2013) and alignments were trimmed to remove poorly aligned and low-coverage regions as previously described (Johnston et al., 2026a), with Nobecovirus (NC_009021) as an outgroup. Maximum-likelihood phylogenies were inferred and visualized as previously described (Johnston et al., 2026a).

### 2.7. Comparative sequence analysis

Complete genomes representing the *P. nathusii*-associated merbecovirus clade were aligned using the MAFFT alignment plugin implemented in Geneious Prime 2026.0.1 with default parameters. Pairwise nucleotide identities were calculated from the resulting whole-genome alignment. Spike genes were subsequently extracted and translated using Geneious Prime. Spike nucleotide and amino acid sequences were aligned separately using the MAFFT plugin with default parameters, and pairwise sequence identities and nucleotide and amino acid substitutions were determined from the resulting alignments.

## 3. Results

### 3.1. Evaluation and optimization of RT-qPCR assays

A manually designed assay targeting the ORF1ab in the bat-associated merbecovirus clade was generated with two different forward primers, sharing a common reverse primer and probe. In addition, nine primer sets were designed using varVAMP (**Figure S1**), primarily targeting ORF1ab, with one set targeting ORF8b. Several of the varVAMP-designed primer sets targeted overlapping regions but differed in their degree of primer and probe degeneracy. Primer sets with higher levels of degeneracy were prioritized, and six varVAMP-designed primer sets were selected for experimental evaluation. Together with the two manually designed primer combinations, a total of eight assays were evaluated (**Table 1**).

The eight assays were initially tested on a merbecovirus reference sample (**Figure S2 A**, **Table S3**). None of the assays generated amplification in the non-template control. Five of the eight assays produced Ct values below 33. Bat-MERS, Bat-MERS_alt, and bat-mers_17 assays yielded Ct values of 31.51, 32.56, and 31.55, respectively, and showed the most robust amplification curves. As these assays targeted the same region within ORF1ab, different combinations of their forward primers, reverse primers, and probes were subsequently evaluated.

All primer-probe combinations generated amplification signals; however, the combination of Bat-MERS-F, bat-mers_17_R, and bat-mers_17_Pr, hereafter termed bat-MERS-F_17Pr-R, produced the lowest Ct value while maintaining robust amplification profile and was therefore selected for further evaluation (**Figure S2 B, Table S4**). To optimize the RT-qPCR conditions, three one-step RT-PCR kits were compared: Qiagen OneStep RT-PCR, AgPath-ID One-Step RT-PCR, and SuperScript III One-Step RT-PCR. The Qiagen OneStep RT-PCR kit produced the most robust amplification profile and was selected for subsequent analyses (**Figure S3**). The optimized bat-MERS-F_17Pr-R assay using the Qiagen OneStep RT-PCR kit was subsequently evaluated against a viral panel comprising bat-associated merbecoviruses, hedgehog-associated merbecoviruses, bat alphacoronaviruses, a bat sarbecovirus, and a hedgehog coronavirus (**Table S2**). The assay detected all bat-associated merbecoviruses included in the panel, with Ct values ranging from 23.38 to 35.92. A hedgehog-associated merbecovirus (ErCoV/38162-6/E.eur/IT/2022) was also detected, albeit at a high Ct-value of 39.50. Among the non-target coronaviruses, weak amplification was observed for one alphacoronavirus (BtCoV/24094-3/P.kuh/IT/2023; Ct 42.96), while the remaining non-target viruses were not detected. The non-template control remained negative.

### 3.2. *In silico* primer analysis

The bat-MERS-F_17Pr-R primer set was evaluated using the primer-check workflow. Screening against the *Merbecovirus* database predicted detection of 898 of 941 sequences (95.4%) using the defined on-target criteria. Applying the less stringent off-target criteria identified one additional potentially detectable sequence. Of the 42 sequences not captured under the on-target criteria, 41 belonged to *Betacoronavirus tylonycteridis* (HKU4). No potential amplification was predicted when the assay was screened against the *Coronaviridae* database excluding *Merbecovirus*, indicating high predicted specificity for merbecoviruses.

### 3.3. Screening for merbecoviruses

Fecal samples from 33 bats collected across Denmark through the 2025 national surveillance program were screened for merbecoviruses using the bat-MERS-F_17Pr-R assay (**Figure 1** and **Table 2**). Two of 33 samples (6.1%) samples tested positive: one from a Nathusius’ pipistrelle (*P. nathusii*; 2025-6602; Ct 32.4) and one from a brown long-eared bat (*P. auratus*; 2025-6604; Ct 31.61). Both RT-qPCR positive samples were subsequently subjected to mNGS for genomic characterization.

**Figure 1.**
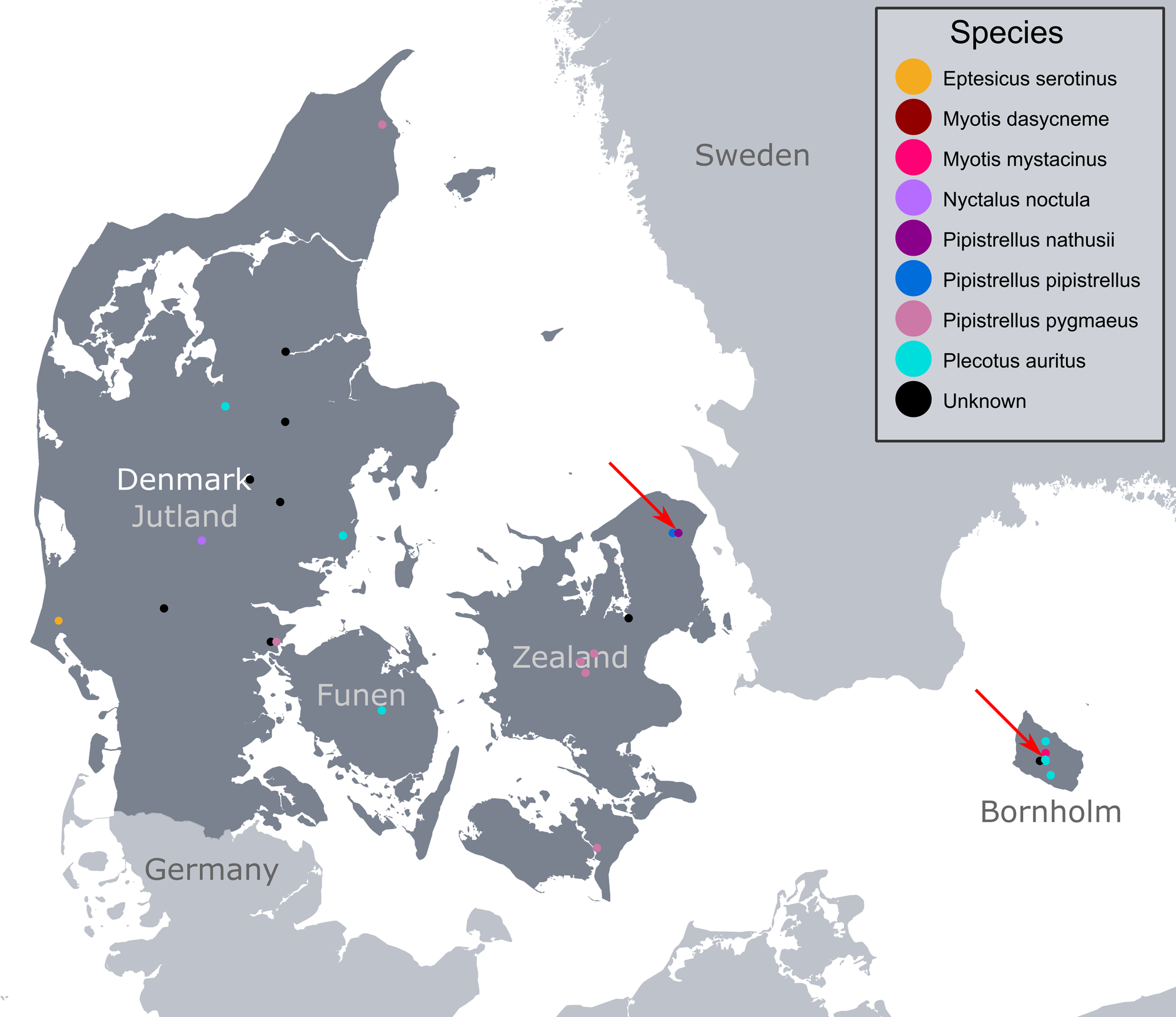
Geographic distribution of bat sampling locations across Denmark. Points represent unique species-location combinations after filtering for duplicate records. Where multiple species were recorded at the same location, jitter was applied for visualization purposes. *Eptesicus serotinus* (yellow), *Myotis dasycneme* (dark red), *Myotis mystacinus* (hot pink), *Nyctalus noctula* (lavender), *Pipistrellus nathusii* (dark magenta), *Pipistrellus pipistrellus* (cobalt blue), *Pipistrellus pygmaeus* (dusty rose), *Plecotus auritus* (cyan), and unknown species (black). The red arrows indicate the locations of the merbecovirus positive samples.

**Table 2:**
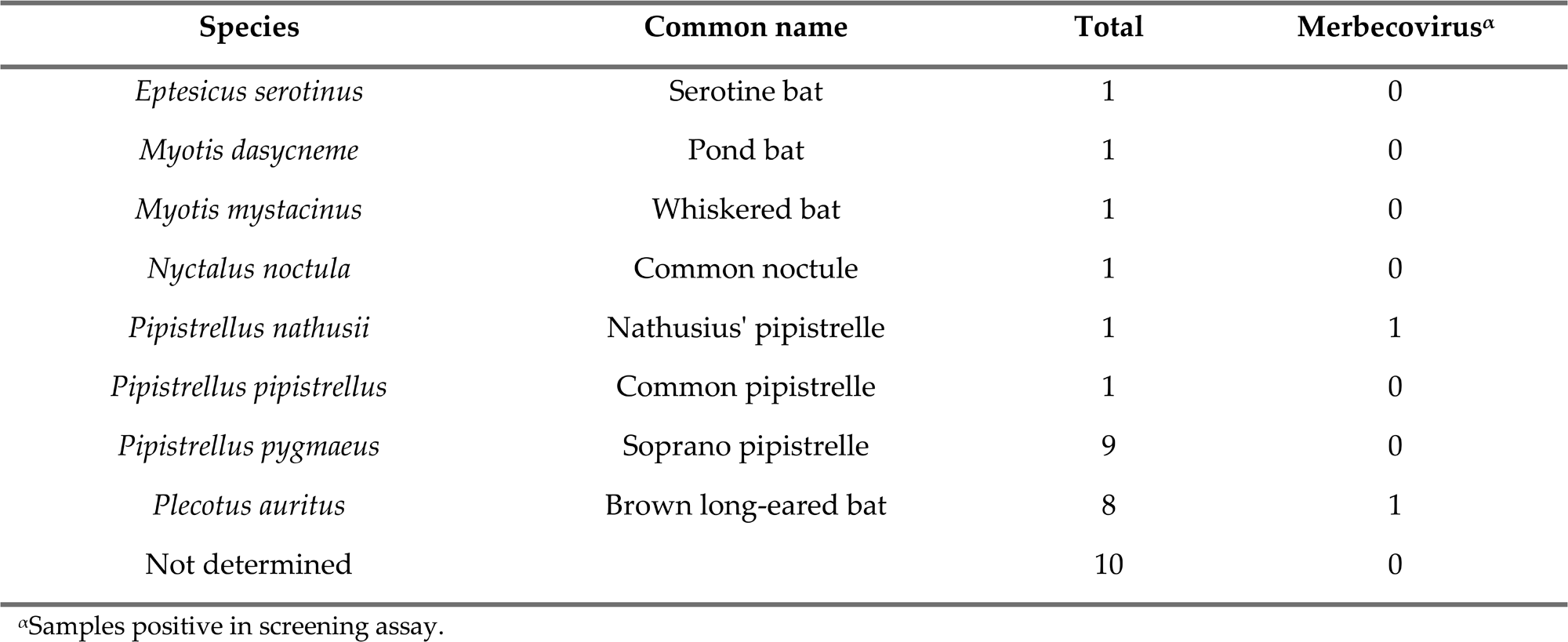
Sampling overview.

### 3.4. Genome recovery and phylogenetic analysis

A complete merbecovirus genome was recovered from the *P. nathusii* sample. A total of 229,168 reads mapped to the recovered genome, resulting in 99.97 % genome coverage at ≥10× depth with no ambiguous nucleotides. Only 440 merbecovirus reads were obtained from the *P. auritus* sample, precluding *de novo* genome assembly. However, reference mapping enabled the generation of a low-coverage consensus sequence, which indicated that the virus was closely related to the previously identified Danish *P. auritus*-associated lineage (**Figure S4**).

A global phylogenetic analysis based on alignment-free pairwise distances confirmed that the complete genome recovered *P. nathusii* belonged to the subgenus *Merbecovirus* (**Figure 2**). A subsequent maximum-likelihood phylogenetic analysis based on a refined sequence alignment showed that the *P. nathusii*-associated virus clustered with merbecoviruses previously identified in *P. nathusii* from Russia (MOW15-22) and the Netherlands (PnNL2018B), forming a distinct clade (**Figure 3**). This clade clustered within a larger lineage comprising merbecoviruses identified predominately in vespertilionid bats, including *Eptesicus*, *Hypsugo*, *Ia*, *Myotis*, *Pipistrellus, Plecotus*, and *Vespertilio* species from Western Europe and East Asia. One related sequence originated from *Rhinolophus ferrumquinum* in South Korea, representing a host association distinct from the otherwise vespertilionid-dominated lineage.

**Figure 2.**
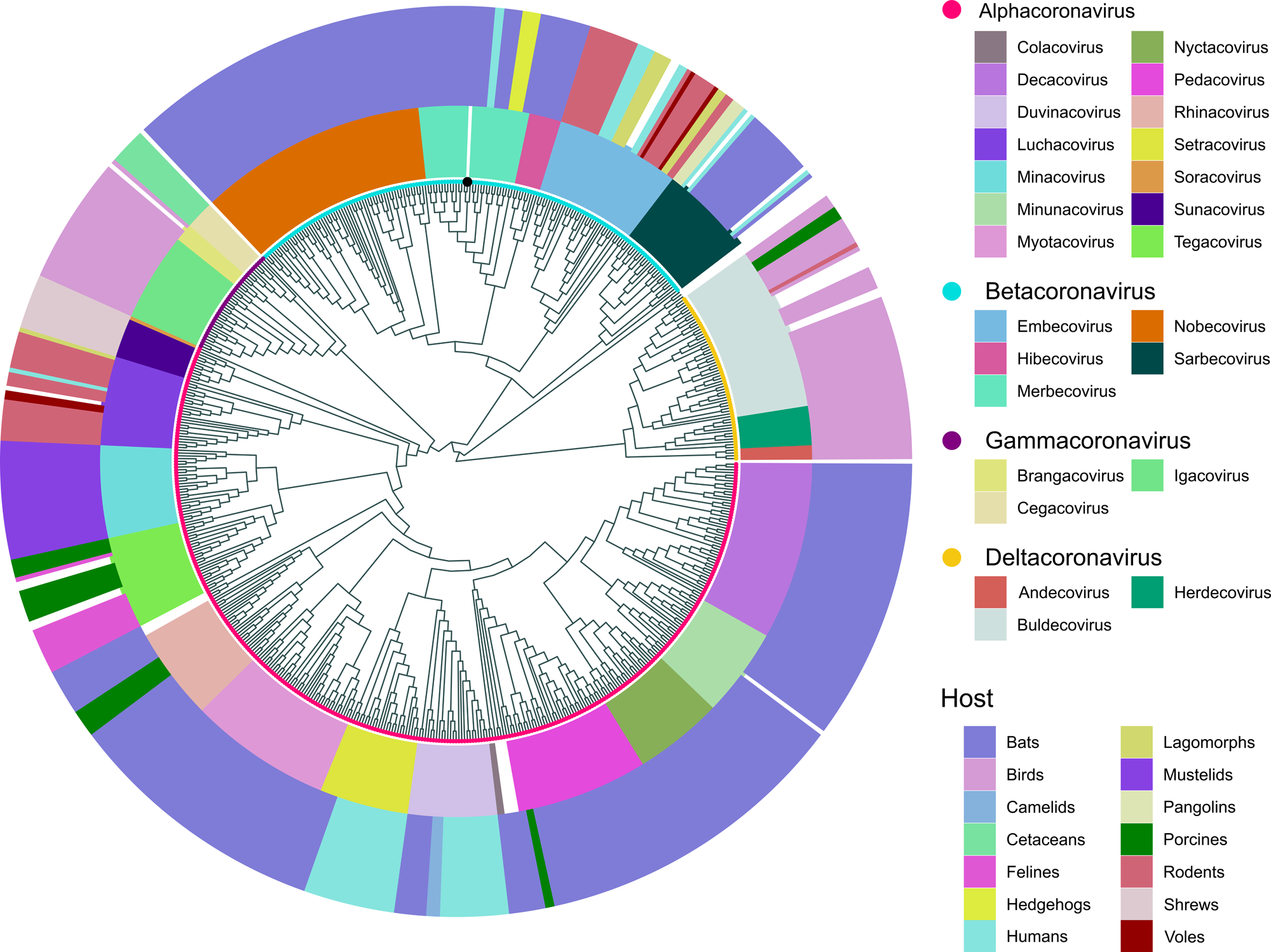
Alignment-free phylogeny of global coronavirus full-length genome diversity (n = 620), including the novel coronavirus genome. The inner ring denotes viral genus (novel genome denoted with a black circle), while the middle and outer rings denote viral subgenus and broader host groups, respectively.

**Figure 3.**
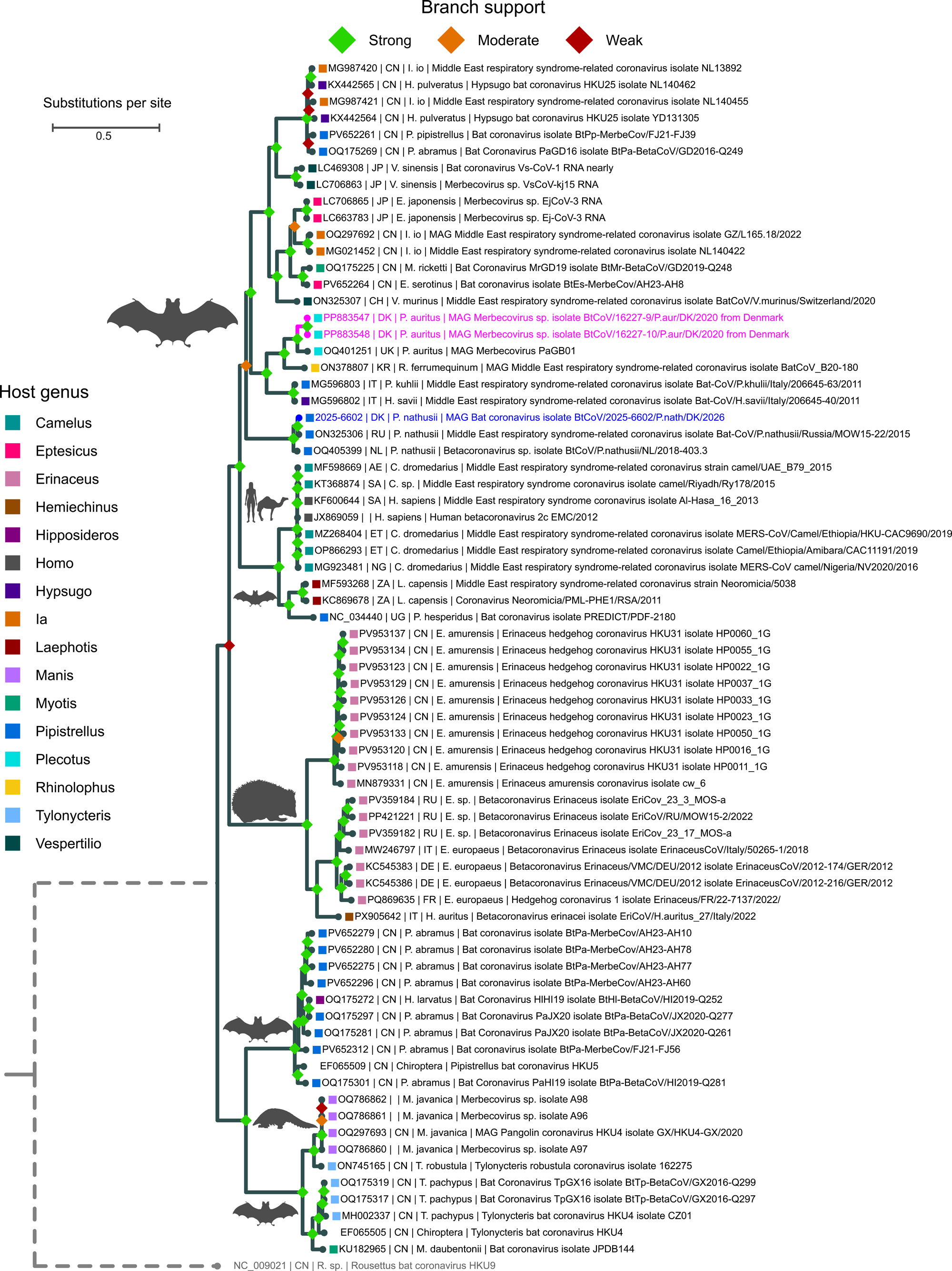
Rooted maximum-likelihood tree of full-length nucleotide sequences of merbecoviruses with the novel virus obtained from a Danish *Pipistrellus nathusii* in blue, previously reported Danish sequences in magenta, and the outgroup in grey. Branch support is categorized as strong (SH-aLRT ≥ 80 and UFBoot ≥ 95), moderate (SH-aLRT ≥ 60 and UFBoot ≥ 60), and weak (below these thresholds). Phylogenetic inference was performed in IQ-TREE using the GTR+F+I+G4 substitution model with 1,000 replicates for both SH-aLRT and UFBoot. Solid branches are proportional to the scale bar (substitutions per site), while dashed branches have been truncated for visualization purposes.

### 3.5. Whole-genome and spike gene comparison

Comparative alignment of the complete genomes within the *P. nathusii*-associated clade showed that the Danish virus 2025-6602 was most closely related to the Russian MOW15-22 strain, sharing 98.67% nucleotide identity across the complete genome. PnNL2018B shared 95.76% and 95.52% genome-wide identity with MOW15-22 and 2025-6602, respectively (**Figure 4**). The greatest sequence divergence among the three genomes was observed within the spike gene.

**Figure 4.**
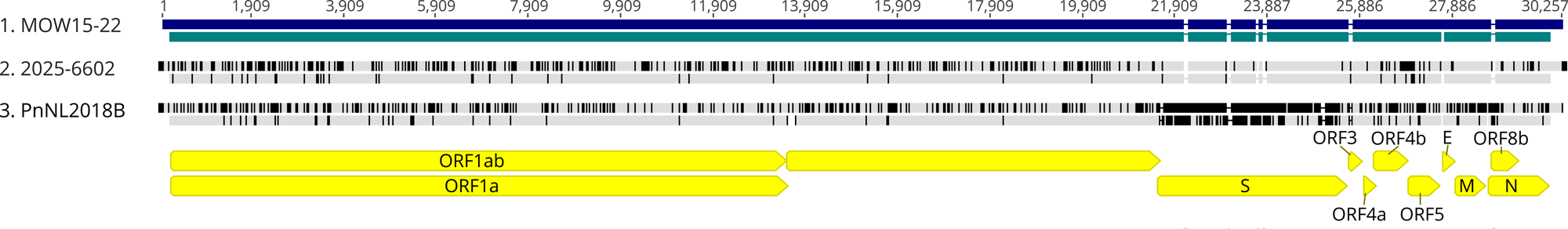
Comparison of full-length sequences from the *Pipistrellus nathusii*-associated merbecovirus clade, comprising the Russian MOW15-22, the Dutch PnNL2018B, and the Danish 2025-6602 viruses. For each sequence, the upper bar represents the nucleotide sequence and the lower bar the translated amino acid sequence. The reference sequence MOW15-22 is shown in dark blue (nucleotide) and petrol (amino acid). Black vertical lines indicate differences relative to the reference sequence.

Pairwise comparison of the spike genes at the nucleotide and amino acid levels showed that 2025-6602 differed from MOW15-22 by only 6 nucleotide substitutions (**Figure S5**). Two substitutions were non-synonymous, resulting in amino acid changes A27V (C80T) and P488S (C1468T), while four were synonymous (C1634T, A2068G, T2341C, and T3970C). Of the two amino acid substitutions, only P488S was located within the RBD. Accordingly, the spike proteins of 2025-6602 and MOW15-22 shared 99.85% amino acid identity. In contrast, the PnNL2018B spike protein was considerably more divergent, sharing only 85.59% amino acid identity with both viruses. Together with the whole-genome phylogenetic analysis, these findings support assignment of the Danish *P. Nathusii*-associated virus to the MOW clade.

## 4. Discussion

In this study, we developed and validated a novel broad-range RT-qPCR assay for the detection of bat-associated merbecoviruses and demonstrated its application in bat surveillance. Although the assay was initially designed to target a phylogenetically defined group of bat-associated merbecoviruses, both experimental validation and *in silico* analysis indicated broader reactivity within the subgenus *Merbecovirus*. The assay detected all bat-associated merbecoviruses included in the experimental panel, while only weak amplification was observed of a single alphacoronavirus. Application of the assay to samples collected through the Danish national bat surveillance program, confirmed the continued presence of merbecoviruses in *P. auritus* and, importantly, identified a merbecovirus in *P. nathusii* for the first time in Denmark. Recovery of a complete genome from the *P. nathusii* sample enabled detailed phylogenetic characterization and assigned the virus to the MOW clade, closely related to a merbecovirus previously identified in *P. nathusii* in Russia.

*P. nathusii* is known as a long-distance migrant that even may crosses large stretches of open sea (Ahlén et al., 2009). A substantial proportion of the population undertakes seasonal movements between summer breeding areas in northeastern-eastern Europe and hibernation areas in western and southwestern Europe (Kruszynski et al., 2021; Pētersons, 2004; Russ, 2020). Migration distances of some marked individuals show examples of migration movements longer than 2000 km from e.g. Finland to United Kingdom, from Latvia to northern Spain and from western Russia to southern France (Alcalde et al., 2021; Russ, 2020). Individuals may cover more than 100 km in a single night. Consequently, the marked individuals may originate from summer habitats further north or east of the first trapping sites.

Nursery roosts are typically located in crevices in trees and in buildings (Baagøe, 2007; Russ, 2020). During the late summer and autumn migration, mating occurs in temporary roosts in trees, buildings or bird and bat boxes. Males apparently make stops at stages in migration areas establishing temporary territories and use acoustic displays to attract migrating females. Harems comprising up to 9 females at a time can be present in the confined spaces of the mating roosts creating ideal conditions for virus transmission between individual from vast ‘upland’ areas of the species summer range (Gerell-Lundberg and Gerell, 1994). Such transient aggregations may bring together individuals originating from geographically separated summer populations, potentially providing opportunities for virus transmission and exchange between otherwise spatially separated bat populations. Together with the long-distance migratory behavior of *P. nathusii*, this ecology may contribute to the dispersal of associated viruses across large geographic areas.

Whole-genome phylogenetic analysis grouped the Danish *P. nathusii*-associated virus with the two previously reported MOW-clade viruses from *P. nathusii* in Russia and the Netherlands, forming a distinct lineage. Comparative genome analysis demonstrated that the Danish virus was considerably more similar to the Russian MOW15-22 genome, than to the Dutch PnNL2018B virus, with the greatest divergence among the three genomes occurring in the spike gene. Comparative spike analysis revealed only two amino acid substitutions between the Danish virus and MOW 15-22, only one of which was located in the RBD.

Recent functional studies have demonstrated that MOW-clade merbecoviruses can utilize ACE2 for cell entry, but do not utilize human ACE2 (Ma et al., 2025). Interestingly, MOW15-22 carries a five-amino acid insertion within the RBD that enables more extensive interactions with ACE2 than the closely related PnNL2018B virus, allowing entry via several non-bat mammalian ACE2 orthologs, while retaining a relatively narrow overall host receptor range (Ma et al., 2025). Although our comparative spike analysis demonstrated near-identical spike proteins between the Danish virus and MOW15-22, suggesting similar receptor usage. However, receptor usage and host range cannot be inferred from sequence similarity alone, and experimental studies will be required to determine the receptor-binding and entry properties of the Danish virus.

The migratory and mating behavior of *P. nathusii* provides a plausible explanation for the high genetic similarity between the Danish virus and the Russian MOW15-22 virus. Extensive movements and mixing of individuals from geographically separated summer populations during migration and at mating roosts may facilitate virus transmission across the distribution range of the species, and thereby limit geographic structuring of associated viruses. However, the small number of available *P. nathusii*-associated merbecovirus genomes currently precludes firm conclusions regarding the relationship between host migration and virus dispersal.

In contrast, *P. auritus* is considered highly sedentary, with limited exchange between local populations, as discussed by Johnston et al., 2025. Detection of another merbecovirus-positive *P. auritus* sample from Bornholm approximately five years after the initial detection is consistent with continued circulation of the previously described *P. auritus-*associated lineage. Although only 440 merbecovirus reads were recovered, precluding *de novo* assembly, reference-based mapping generated a partial, low-coverage consensus sequence consistent with assignment to the same lineage. The limited sequence coverage precludes a robust assessment of genetic divergence from the previously characterized viruses. This lineage belongs to the broader HKU25 clade but appears to be distinguished by a characteristic deletion within the RBD. Continued surveillance of *P. auritus* populations will be important to determine the prevalence, persistence and evolutionary dynamics of this lineage.

Metagenomic sequencing provides the genomic information required for virus discovery, phylogenetic analysis, and characterization of viral diversity. However, successful genome reconstruction depends on several factors, including viral abundance, sequencing depth, and the assembly strategy applied. These limitations were illustrated in the present study by the recovery of a complete genome from the *P. nathusii* sample, whereas only a partial, low-coverage consensus sequence could be generated from the *P. auritus* sample despite successful detection by RT-qPCR. As discussed previously (Johnston et al., 2026a, 2025), both *de novo* assembly and reference-guided approaches have inherent limitations, including fragmented assemblies resulting from uneven sequencing depth and potential reference bias, particularly in highly variable genomic regions such as the spike gene.

Targeted RT-qPCR and metagenomic sequencing therefore provide complementary approaches for wildlife virus surveillance. RT-qPCR provides a rapid, sensitive, and cost-effective method for screening large numbers of samples, while probe-based detection enables immediate identification of the target virus group and prioritization of positive samples for downstream sequencing. Metagenomic sequencing can subsequently provide genomic information for phylogenetic and evolutionary analyses when sufficient viral material is present. Combining targeted molecular screening with mNGS thus provides an efficient surveillance strategy that couples high-throughput detection with genomic characterization of circulating viruses.

In conclusion, we developed a broadly inclusive RT-qPCR assay for bat-associated merbecoviruses and demonstrated its utility for wildlife surveillance. Application of the assay provided evidence of continued circulation of the previously identified Danish *P. auritus*-associated lineage and enabled the detection and genomic characterization of a MOW-clade merbecovirus in *P. nathusii* for the first time in Denmark Together, these findings expand our understanding of merbecovirus diversity in Danish bats and provide a practical molecular tool for future surveillance and genomic characterization of this diverse group of bat-associated coronaviruses.

## Supplementary

**Table S1**: Bat associated merbecovirus clade dataset

**Table S2:** Virus panel for assay validation

**Table S3:** Assay evaluation

**Table S4:** Combination evaluation

**Table S5**: Bat sample collection details

**Table S6:** Coronavirus representative dataset

**Table S7:** Merbecovirus representative dataset

**Figure S1.** Placement of designed primers on the Danish merbecovirus BtCoV/16227-9/P.aur/DK/2020.

**Figure S2.** Amplification curves for evaluated primer sets. A) Bat-MERS (lavender), Bat-MERS_alt (hot pink), bat-mers_16 (golden yellow), bat-mers_17 (cyan), bat-mers_18 (dark red), bat-mers_19a (cobalt blue), bat-mers_19b (dark magenta), and bat-mers_28 (sea green). B) Bat-MERS (lavender), Bat-MERS_alt (hot pink), bat-mers_17 (cyan), Bat-MERS_pr17 (dark teal), bat-mers-17F-Pr_MERS-R (dusty rose), bat-mers-17F_MERS-Pr-R (burnt orange), and bat-MERS-F_17Pr-R (light blue). Merbecovirus positive and negative control as solid and dashed lines, respectively. Curves are shown for cycles 10-45.

**Figure S3.** Amplification curves for selected primer set bat-MERS-F_17Pr-R. A) Baseline Subtracted Curve Fit plot. B) No Baseline Subtraction plot. AgPath (dark red), Qiagen (golden yellow) and SuperScript III (dark magenta) kits. Merbecovirus positive and negative control as solid and dashed lines, respectively. Curves are shown for cycles 10-45.

**Figure S4.** Alignment of the low-coverage consensus sequence of the novel *Plecotus auritus* merbecovirus (2025-6604) against the previously identified Danish *P. auritus*-associated merbecovirus BtCoV/16227-9/P.aur/DK/2020 (PP883547.1). Matching positions are shown as light grey bars, single nucleotide polymorphisms (SNPs) as magenta bars, ambiguous nucleotides as dark grey bars, and regions without sequence coverage as black horizontal lines.

**Figure S5.** Comparison of spike gene sequences from the *Pipistrellus nathusii*-associated merbecovirus clade, comprising the Russian MOW15-22, the Dutch PnNL2018B, and the Danish 2025-6602 viruses. For each sequence, the upper bar represents the nucleotide sequence and the lower bar the translated amino acid sequence. The reference sequence MOW15-22 is shown in dark blue (nucleotide) and petrol (amino acid). Black vertical lines indicate differences relative to the reference sequence.

## Declarations

### Ethical approval

Ethical review and approval were waived for this study. Samples from live bats consisted of fecal material collected non-invasively under dispensations MST-850-00116 and SN 302-009 issued by the Danish Environmental Protection Agency. Samples included through the passive surveillance were obtained exclusively from animals that were already deceased.

### Consent for publication

Not applicable.

### Availability of data and materials

The datasets used and/or analyzed during the current study are available from the corresponding author on reasonable request.

Sequencing data are available at NCBI BioProject with accession no. PRJNA1536352. The complete merbecovirus genome recovered from *P. Nathusii* has been deposited in GenBank under accession no. QB139754.

## Author’s contribution

**CMJ:** Conceptualization, Data curation, Formal analysis, Investigation, Methodology, Project administration, Visualization, Writing – original draft, Writing – review and editing. **AM:** Resources, Writing – review and editing. **DL:** Resources, Writing – review and editing. **ELM:** Methodology, Writing – review and editing. **HJB:** Investigation, Resources, Writing – review and editing. **ME:** Investigation, Resources, Writing – review and editing. **ETF:** Investigation, Resources, Writing – review and editing. **LL:** Funding acquisition, Supervision, Writing – review and editing. **TBR:** Conceptualization, Formal analysis, Funding acquisition, Project administration, Supervision, Writing – review and editing.

## Acknowledgements

We thank Jani Christiansen, Rasmus Oskar Rask Hansen, and Fie Fisker Brønnum Christensen for their invaluable technical assistance during this study. We also thank Anne Sofie Vedsted Hammer, Tim Kåre Jensen, and Michelle Lauge Quaade for their contributions to the examinations of bat carcasses.

## References

Ahlén, I., Baagøe, H.J., Bach, L., 2009. Behavior of Scandinavian Bats during Migration and Foraging at Sea. J. Mammal. 90, 1318–1323. 10.1644/09-MAMM-S-223R.1

Alcalde, J.T., Jiménez, M., Brila, I., Vintulis, V., Voigt, C.C., Pētersons, G., 2021. Transcontinental 2200 km migration of a Nathusius’ pipistrelle ( Pipistrellus nathusii ) across Europe. Mammalia 85, 161–163. 10.1515/mammalia-2020-0069

Annan, A., Baldwin, H.J., Corman, V.M., Klose, S.M., Owusu, M., Nkrumah, E.E., Badu, E.K., Anti, P., Agbenyega, O., Meyer, B., Oppong, S., Sarkodie, Y.A., Kalko, E.K.V., Lina, P.H.C., Godlevska, E. V., Reusken, C., Seebens, A., Gloza-Rausch, F., Vallo, P., Tschapka, M., Drosten, C., Drexler, J.F., 2013. Human Betacoronavirus 2c EMC/2012–related Viruses in Bats, Ghana and Europe. Emerg. Infect. Dis. 19, 456–459. 10.3201/eid1903.121503

Anthony, S.J., Gilardi, K., Menachery, V.D., Goldstein, T., Ssebide, B., Mbabazi, R., Navarrete-Macias, I., Liang, E., Wells, H., Hicks, A., Petrosov, A., Byarugaba, D.K., Debbink, K., Dinnon, K.H., Scobey, T., Randell, S.H., Yount, B.L., Cranfield, M., Johnson, C.K., Baric, R.S., Lipkin, W.I., Mazet, J.A.K., 2017. Further Evidence for Bats as the Evolutionary Source of Middle East Respiratory Syndrome Coronavirus. MBio 8. 10.1128/mBio.00373-17

Anthony, S.J., Ojeda-Flores, R., Rico-Chávez, O., Navarrete-Macias, I., Zambrana-Torrelio, C.M., Rostal, M.K., Epstein, J.H., Tipps, T., Liang, E., Sanchez-Leon, M., Sotomayor-Bonilla, J., Aguirre, A.A., Ávila-Flores, R., Medellín, R.A., Goldstein, T., Suzán, G., Daszak, P., Lipkin, W.I., 2013. Coronaviruses in bats from Mexico. J. Gen. Virol. 94, 1028–1038. 10.1099/vir.0.049759-0

Baagøe, H.J., 2007. Flagermus, Chiroptera, in: Baagøe, H.J., Jensen, T.S. (Eds.), Dansk Pattedyratlas. Gyldendal, pp. 38–98.

Camacho, C., Coulouris, G., Avagyan, V., Ma, N., Papadopoulos, J., Bealer, K., Madden, T.L., 2009. BLAST+: architecture and applications. BMC Bioinformatics 10, 421. 10.1186/1471-2105-10-421

Catanzaro, N.J., Wu, Z., Fan, C., Jefferson, V., Abdelgadir, A., Schäfer, A., Yount, B.L., Bjorkman, P.J., Baric, R., Letko, M., 2025. ACE2 from Pipistrellus abramus bats is a receptor for HKU5 coronaviruses. Nat. Commun. 16, 4932. 10.1038/s41467-025-60286-3

Chen, J., Yang, X., Si, H., Gong, Q., Que, T., Li, J., Li, Y., Wu, C., Zhang, W., Chen, Y., Luo, Y., Zhu, Y., Li, B., Luo, D., Hu, B., Lin, H., Jiang, R., Jiang, T., Li, Q., Liu, M., Xie, S., Su, J., Zheng, X., Li, A., Yao, Y., Yang, Y., Chen, P., Wu, A., He, M., Lin, X., Tong, Y., Hu, Y., Shi, Z.-L., Zhou, P., 2023. A bat MERS-like coronavirus circulates in pangolins and utilizes human DPP4 and host proteases for cell entry. Cell 186, 850–863.e16. 10.1016/j.cell.2023.01.019

Chen, J., Zhang, W., Li, Y., Liu, C., Dong, T., Chen, H., Wu, C., Su, J., Li, B., Zhang, W., Hu, B., Jia, J., Ma, C.-B., Zhu, Y., He, X., Li, A., Pan, K., Lin, H., Guo, Z., Li, C., Zhang, L., Yan, H., Zhou, P., Peng, W., Shi, Z.-L., 2025. Bat-infecting merbecovirus HKU5-CoV lineage 2 can use human ACE2 as a cell entry receptor. Cell 188, 1729–1742.e16. 10.1016/j.cell.2025.01.042

Corman, V.M., Ithete, N.L., Richards, L.R., Schoeman, M.C., Preiser, W., Drosten, C., Drexler, J.F., 2014. Rooting the Phylogenetic Tree of Middle East Respiratory Syndrome Coronavirus by Characterization of a Conspecific Virus from an African Bat. J. Virol. 88, 11297–11303. 10.1128/JVI.01498-14

Corman, V.M., Muth, D., Niemeyer, D., Drosten, C., 2018. Hosts and Sources of Endemic Human Coronaviruses. pp. 163–188. 10.1016/bs.aivir.2018.01.001

de Groot, R.J., Baker, S.C., Baric, R., Enjuanes, L., Gorbalenya, A.E., Holmes, K. V., Perlman, S., Poon, L., Rottier, P.J.M., Talbot, P.J., Woo, P.C.Y., Ziebuhr, J., 2012. Family: Coronaviridae, in: Virus Taxonomy, Ninth Report of the International Committee on Taxonomy of Viruses. Elsevier, pp. 806–828. 10.1016/B978-0-12-384684-6.00068-9

De Sabato, L., Di Bartolo, I., De Marco, M.A., Moreno, A., Lelli, D., Cotti, C., Delogu, M., Vaccari, G., 2020. Can Coronaviruses Steal Genes from the Host as Evidenced in Western European Hedgehogs by EriCoV Genetic Characterization? Viruses 12, 1471. 10.3390/v12121471

Fuchs, J., Kleine, J., Schemmerer, M., Kreibich, J., Maier, W., Battur, N., Krannich, T., Sedaghatjoo, S., Jaki, L., Maks, A., Boehm, C., Wilhelm, C., Schulze, J., Mache, C., Berger, E., Panajotov, J., Arnold, L., Grüning, B., Bauswein, M., Böttcher, S., Johne, R., Wenzel, J., Hölzer, M., Panning, M., 2025. varVAMP: degenerate primer design for tiled full genome sequencing and qPCR. Nat. Commun. 16, 5067. 10.1038/s41467-025-60175-9

Gerell-Lundberg, K., Gerell, R., 1994. The mating behaviour of the pipistrelle and the Nathusius’ pipistrelle (Chiroptera)-a comparison. FOLIA Zool. 43, 315.

Haagmans, B.L., Al Dhahiry, S.H.S., Reusken, C.B.E.M., Raj, V.S., Galiano, M., Myers, R., Godeke, G.-J., Jonges, M., Farag, E., Diab, A., Ghobashy, H., Alhajri, F., Al-Thani, M., Al-Marri, S.A., Al Romaihi, H.E., Al Khal, A., Bermingham, A., Osterhaus, A.D.M.E., AlHajri, M.M., Koopmans, M.P.G., 2014. Middle East respiratory syndrome coronavirus in dromedary camels: an outbreak investigation. Lancet Infect. Dis. 14, 140–145. 10.1016/S1473-3099(13)70690-X

He, W.-T., Hou, X., Zhao, J., Sun, J., He, H., Si, W., Wang, J., Jiang, Z., Yan, Z., Xing, G., Lu, M., Suchard, M.A., Ji, X., Gong, W., He, B., Li, J., Lemey, P., Guo, D., Tu, C., Holmes, E.C., Shi, M., Su, S., 2022. Virome characterization of game animals in China reveals a spectrum of emerging pathogens. Cell 185, 1117–1129.e8. 10.1016/j.cell.2022.02.014

Hu, B., Ge, X., Wang, L.-F., Shi, Z., 2015. Bat origin of human coronaviruses. Virol. J. 12, 221. 10.1186/s12985-015-0422-1

Ithete, N.L., Stoffberg, S., Corman, V.M., Cottontail, V.M., Richards, L.R., Schoeman, M.C., Drosten, C., Drexler, J.F., Preiser, W., 2013. Close Relative of Human Middle East Respiratory Syndrome Coronavirus in Bat, South Africa. Emerg. Infect. Dis. 19, 1697–1699. 10.3201/eid1910.130946

Jin, M., Jefferson, V.A., Zhao, Z., Catanzaro, N.J., De Sabato, L., Keller, E.L., Menasche, B.L., Hoffman, C., Castelli, A., Di Bartolo, I., Vaccari, G., Starr, T.N., Seifert, S.N., Castellanos, A.A., Han, B.A., Serebryannyy, L., Spaulding, A.B., Douek, D.C., Moreno, A., Baric, R.S., Wilen, C.B., Rini, J.M., Letko, M., 2025. Aminopeptidase N is a receptor for hedgehog merbecoviruses. bioRxiv. 10.1101/2025.09.03.674079

Johnston, C.M., Gunalan, V., Baagoe, H.J., Elmeros, M., Fjederholt, E.T., Quaade, M.L., Hansen, M.J.S., Lohse, L., Rasmussen, T.B., 2026a. Targeted metagenomic recovery of coronaviruses from wildlife samples. bioRxiv. 10.64898/2026.05.05.722935

Johnston, C.M., Gunalan, V., Baagøe, H.J., Fomsgaard, A.S., Polacek, C., Rasmussen, M., Lohse, L., Rasmussen, T.B., 2025. Characterization of a MERS-related betacoronavirus in Danish brown long-eared bats (Plecotus auritus). Virol. J. 22, 283. 10.1186/s12985-025-02883-8

Johnston, C.M., Mingoli, A., Hoegh, A.M., Lohse, L., Rasmussen, T.B., 2026b. Efficient primer design for broad-range virus detection: A semi-automated workflow using sequence clustering and varVAMP. bioRxiv. 10.64898/2026.09.21.753121

Katoh, K., Standley, D.M., 2013. MAFFT Multiple Sequence Alignment Software Version 7: Improvements in Performance and Usability. Mol. Biol. Evol. 30, 772–780. 10.1093/molbev/mst010

Kruszynski, C., Bailey, L.D., Courtiol, A., Bach, L., Bach, P., Göttsche, Matthias, Göttsche, Michael, Hill, R., Lindecke, O., Matthes, H., Pommeranz, H., Popa-Lisseanu, A.G., Seebens-Hoyer, A., Tichomirowa, M., Voigt, C.C., 2021. Identifying migratory pathways of Nathusius’ pipistrelles ( Pipistrellus nathusii ) using stable hydrogen and strontium isotopes. Rapid Commun. Mass Spectrom. 35. 10.1002/rcm.9031

Lau, S.K.P., Li, K.S.M., Tsang, A.K.L., Lam, C.S.F., Ahmed, S., Chen, H., Chan, K.-H., Woo, P.C.Y., Yuen, K.-Y., 2013. Genetic Characterization of Betacoronavirus Lineage C Viruses in Bats Reveals Marked Sequence Divergence in the Spike Protein of Pipistrellus Bat Coronavirus HKU5 in Japanese Pipistrelle: Implications for the Origin of the Novel Middle East Respiratory Sy. J. Virol. 87, 8638–8650. 10.1128/JVI.01055-13

Lazov, C.M., Belsham, G.J., Bøtner, A., Rasmussen, T.B., 2021. Full-Genome Sequences of Alphacoronaviruses and Astroviruses from Myotis and Pipistrelle Bats in Denmark. Viruses 13, 1073. 10.3390/v13061073

Lazov, C.M., Chriél, M., Baagøe, H.J., Fjederholt, E., Deng, Y., Kooi, E., Belsham, G., Bøtner, A., Rasmussen, T.B., 2018. Detection and Characterization of Distinct Alphacoronaviruses in Five Different Bat Species in Denmark. Viruses 10, 486. 10.3390/v10090486

Lelli, D., Papetti, A., Sabelli, C., Rosti, E., Moreno, A., Boniotti, M., 2013. Detection of Coronaviruses in Bats of Various Species in Italy. Viruses 5, 2679–2689. 10.3390/v5112679

Letko, M., Seifert, S.N., Olival, K.J., Plowright, R.K., Munster, V.J., 2020. Bat-borne virus diversity, spillover and emergence. Nat. Rev. Microbiol. 18, 461–471. 10.1038/s41579-020-0394-z

Liu, C., Park, Y.-J., Ma, C.-B., Stuart, C., Gen, R., Sun, Y.-C., Yang, X., Lin, M.-Y., Xiong, Q., Si, J.-Y., Liu, P., Veesler, D., Yan, H., 2025. ACE2 utilization of HKU25 clade MERS-related coronaviruses with broad geographic distribution. bioRxiv. 10.1101/2025.02.19.639017

Luo, C.-M., Wang, N., Yang, X.-L., Liu, H.-Z., Zhang, W., Li, B., Hu, B., Peng, C., Geng, Q.-B., Zhu, G.-J., Li, F., Shi, Z.-L., 2018. Discovery of Novel Bat Coronaviruses in South China That Use the Same Receptor as Middle East Respiratory Syndrome Coronavirus. J. Virol. 92. 10.1128/JVI.00116-18

Ma, C.-B., Liu, C., Park, Y.-J., Tang, J., Chen, J., Xiong, Q., Lee, J., Stewart, C., Asarnow, D., Brown, J., Tortorici, M.A., Yang, X., Sun, Y.-H., Chen, Y.Y.-M., Yu, X., Si, J.J.-Y., Liu, P., Tong, F., Huang, M.-L.M., Li, J., Shi, Z.-L., Deng, Z., Veesler, D., Yan, H., 2025. Multiple independent acquisitions of ACE2 usage in MERS-related coronaviruses. Cell 188, 1693–1710.e18. 10.1016/j.cell.2024.12.031

Ma, C.-B., Liu, C., Xiong, Q., Gu, M., Shi, L., Wang, C., Si, J., Tong, F., Liu, P., Huang, M., Yan, H., 2023. Broad host tropism of ACE2-using MERS-related coronaviruses and determinants restricting viral recognition. Cell Discov. 9, 57. 10.1038/s41421-023-00566-8

Memish, Z.A., Mishra, N., Olival, K.J., Fagbo, S.F., Kapoor, V., Epstein, J.H., Alhakeem, R., Durosinloun, A., Al Asmari, M., Islam, A., Kapoor, A., Briese, T., Daszak, P., Al Rabeeah, A.A., Lipkin, W.I., 2013. Middle East respiratory syndrome coronavirus in bats, Saudi Arabia. Emerg. Infect. Dis. 19, 1819–23. 10.3201/eid1911.131172

Moreno, A., Lelli, D., de Sabato, L., Zaccaria, G., Boni, A., Sozzi, E., Prosperi, A., Lavazza, A., Cella, E., Castrucci, M.R., Ciccozzi, M., Vaccari, G., 2017. Detection and full genome characterization of two beta CoV viruses related to Middle East respiratory syndrome from bats in Italy. Virol. J. 14, 239. 10.1186/s12985-017-0907-1

Nooij, S., Schmitz, D., Vennema, H., Kroneman, A., Koopmans, M.P.G., 2018. Overview of Virus Metagenomic Classification Methods and Their Biological Applications. Front. Microbiol. 9. 10.3389/fmicb.2018.00749

Park, Y.-J., Liu, C., Lee, J., Brown, J.T., Ma, C.-B., Liu, P., Gen, R., Xiong, Q., Zepeda, S.K., Stewart, C., Addetia, A., Craig, C.J., Tortorici, M.A., Alshukairi, A.N., Starr, T.N., Yan, H., Veesler, D., 2025. Molecular basis of convergent evolution of ACE2 receptor utilization among HKU5 coronaviruses. Cell 188, 1711–1728.e21. 10.1016/j.cell.2024.12.032

Pētersons, G., 2004. Seasonal migrations of north-eastern populations of Nathusius’ bat Pipistrellus nathusii (Chiroptera). Myotis 41, 29–56.

Russ, J., 2020. Nathusius’s Pipistrelle Pipistrellus nathusii (Keyserling and Blasius, 1839), in: Hackländer, K., Zachos, F.E. (Eds.), Handbook of the Mammals of Europe. Springer International Publishing, Cham, pp. 1–26. 10.1007/978-3-319-65038-8_68-1

Schlegel, M., Ali, H.S., Stieger, N., Groschup, M.H., Wolf, R., Ulrich, R.G., 2012. Molecular Identification of Small Mammal Species Using Novel Cytochrome b Gene-Derived Degenerated Primers. Biochem. Genet. 50, 440–447. 10.1007/s10528-011-9487-8

Sola, I., Almazan, F., Zúñiga, S., Enjuanes, L., 2015. Continuous and Discontinuous RNA Synthesis in Coronaviruses. Annu. Rev. Virol. 2, 265–288. 10.1146/annurev-virology-100114-055218

Temmam, S., Vongphayloth, K., Baquero, E., Munier, S., Bonomi, M., Regnault, B., Douangboubpha, B., Karami, Y., Chrétien, D., Sanamxay, D., Xayaphet, V., Paphaphanh, P., Lacoste, V., Somlor, S., Lakeomany, K., Phommavanh, N., Pérot, P., Dehan, O., Amara, F., Donati, F., Bigot, T., Nilges, M., Rey, F.A., van der Werf, S., Brey, P.T., Eloit, M., 2022. Bat coronaviruses related to SARS-CoV-2 and infectious for human cells. Nature 604, 330–336. 10.1038/s41586-022-04532-4

van Boheemen, S., de Graaf, M., Lauber, C., Bestebroer, T.M., Raj, V.S., Zaki, A.M., Osterhaus, A.D.M.E., Haagmans, B.L., Gorbalenya, A.E., Snijder, E.J., Fouchier, R.A.M., 2012. Genomic Characterization of a Newly Discovered Coronavirus Associated with Acute Respiratory Distress Syndrome in Humans. Mbio 3. 10.1128/mBio.00473-12

Wang, Q., Qi, J., Yuan, Y., Xuan, Y., Han, P., Wan, Y., Ji, W., Li, Y., Wu, Y., Wang, J., Iwamoto, A., Woo, P.C.Y., Yuen, K.-Y., Yan, J., Lu, G., Gao, G.F., 2014. Bat Origins of MERS-CoV Supported by Bat Coronavirus HKU4 Usage of Human Receptor CD26. Cell Host Microbe 16, 328–337. 10.1016/j.chom.2014.08.009

Woo, P.C.Y.Y., de Groot, R.J., Haagmans, B., Lau, S.K.P., Neuman, B.W., Perlman, S., Sola, I., van der Hoek, L., Wong, A.C.P., Yeh, S.-H., 2023. ICTV Virus Taxonomy Profile: Coronaviridae 2023. J. Gen. Virol. 104. 10.1099/jgv.0.001843

Xiong, Q., Cao, L., Ma, C.-B., Liu, C., Si, J., Liu, P., Gu, M., Wang, C., Shi, L., Tong, F., Huang, M., Li, J., Zhao, C., Shen, C., Chen, Y., Zhao, H., Lan, K., Wang, X., Yan, H., 2022. Close relatives of MERS-CoV in bats use ACE2 as their functional receptors. Nature. 10.1101/2022.01.24.477490

Yang, L., Wu, Z., Ren, X., Yang, F., Zhang, J., He, G., Dong, J., Sun, L., Zhu, Y., Zhang, S., Jin, Q., 2014. MERS–Related Betacoronavirus in Vespertilio superans Bats, China. Emerg. Infect. Dis. 20. 10.3201/eid2007.140318

Yang, Y., Du, L., Liu, C., Wang, L., Ma, C., Tang, J., Baric, R.S., Jiang, S., Li, F., 2014. Receptor usage and cell entry of bat coronavirus HKU4 provide insight into bat-to-human transmission of MERS coronavirus. Proc. Natl. Acad. Sci. 111, 12516–12521. 10.1073/pnas.1405889111

Zielezinski, A., Gudyś, A., Barylski, J., Siminski, K., Rozwalak, P., Dutilh, B.E., Deorowicz, S., 2025. Ultrafast and accurate sequence alignment and clustering of viral genomes. Nat. Methods 22, 1191–1194. 10.1038/s41592-025-02701-7

