## Supplementary figures and images for "Detection of merbecoviruses in Danish bats using a novel broad-range RT-qPCR assay"

### Supplementary figure S01

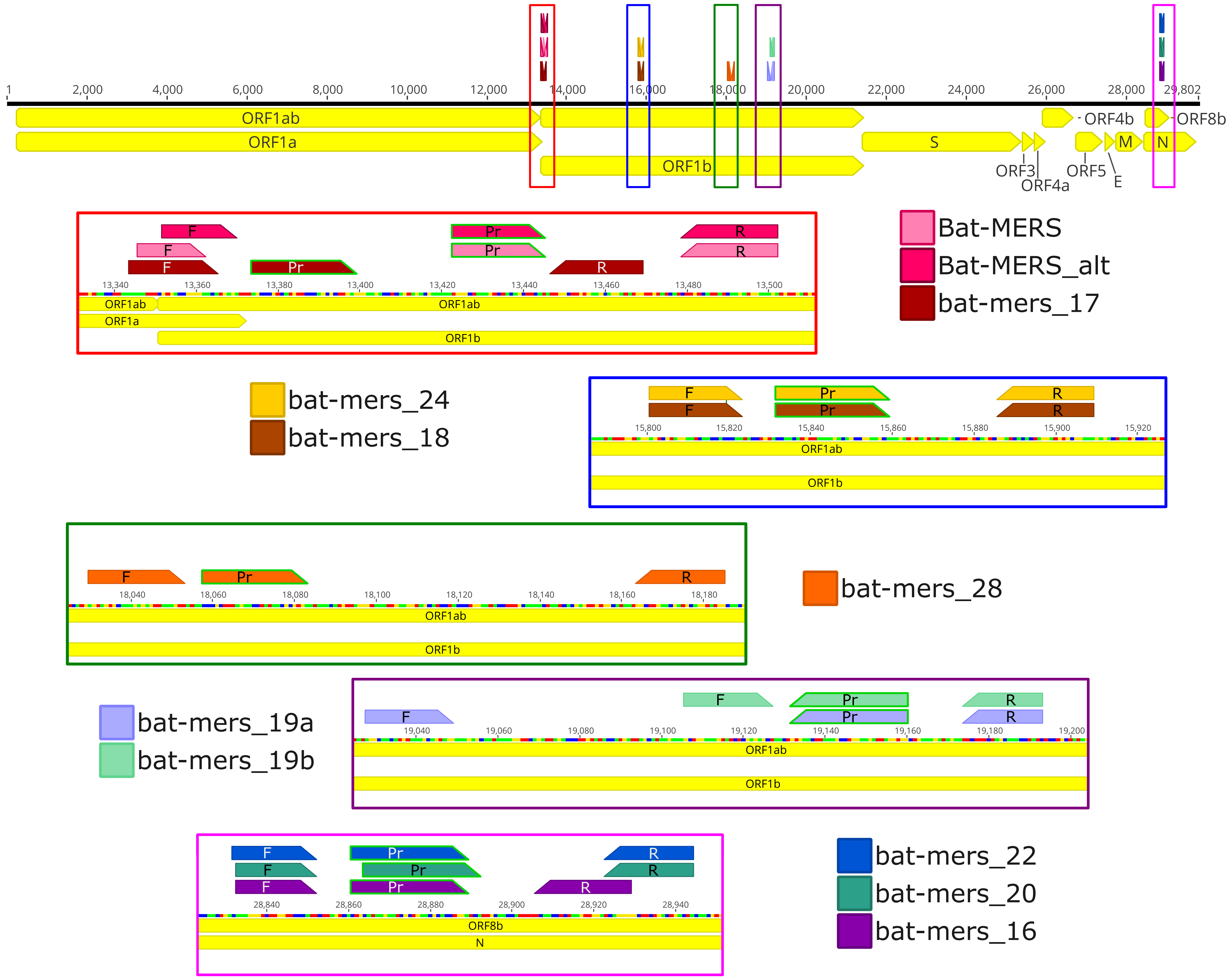

### Supplementary figure S02

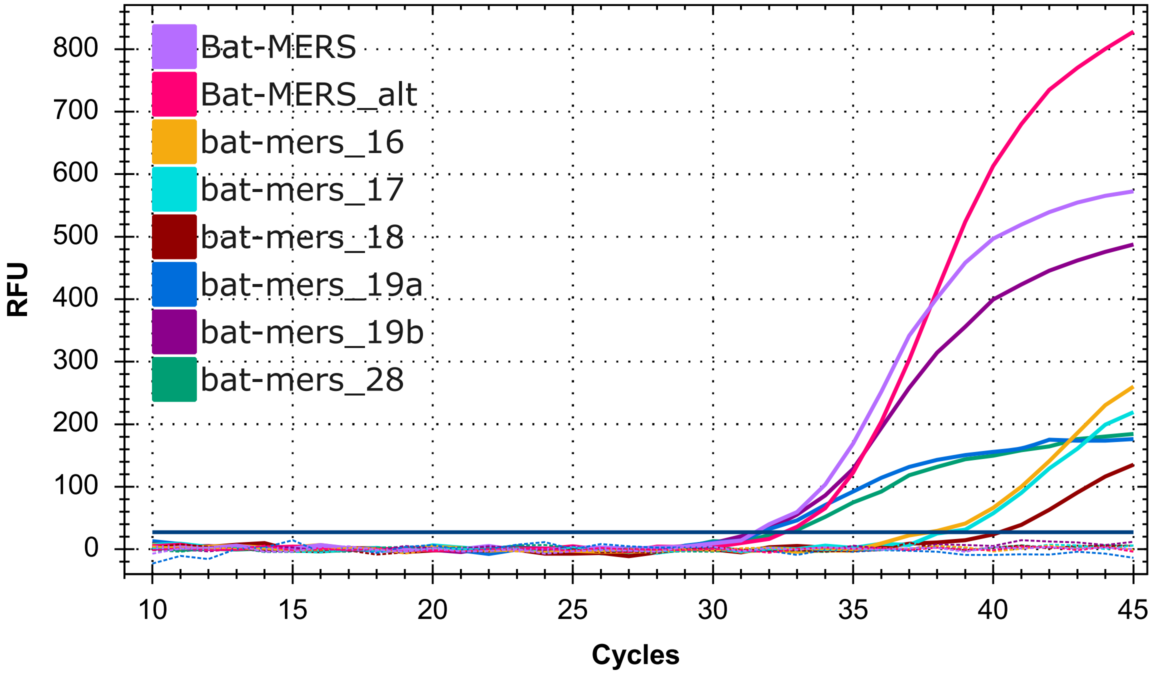

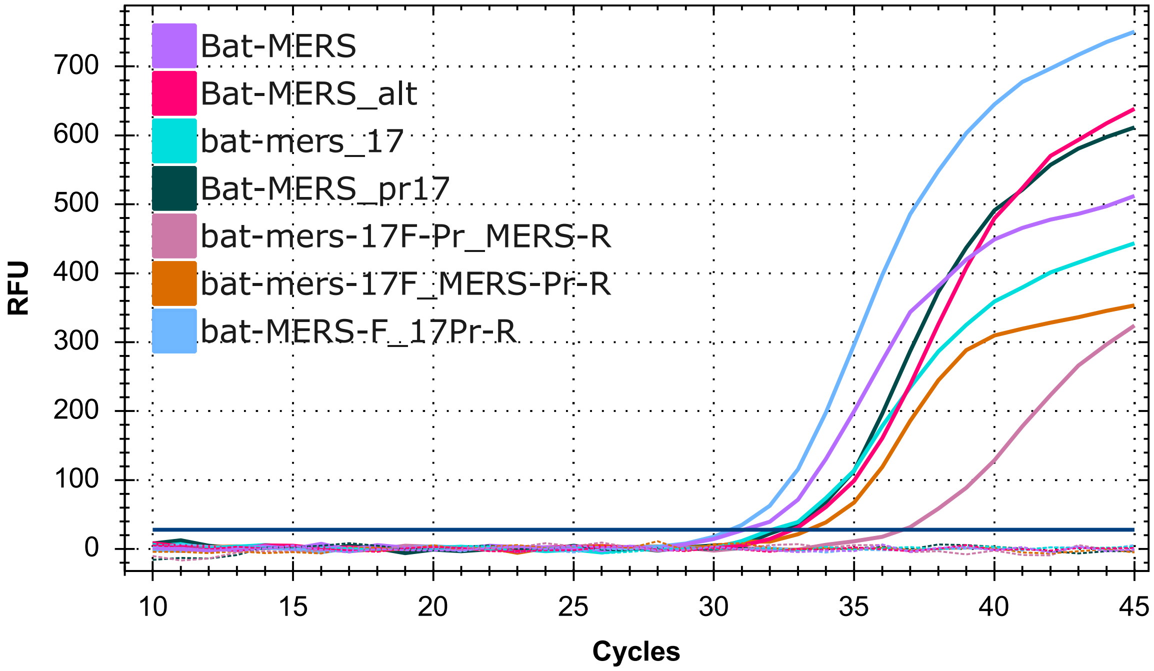

### Supplementary figure S03

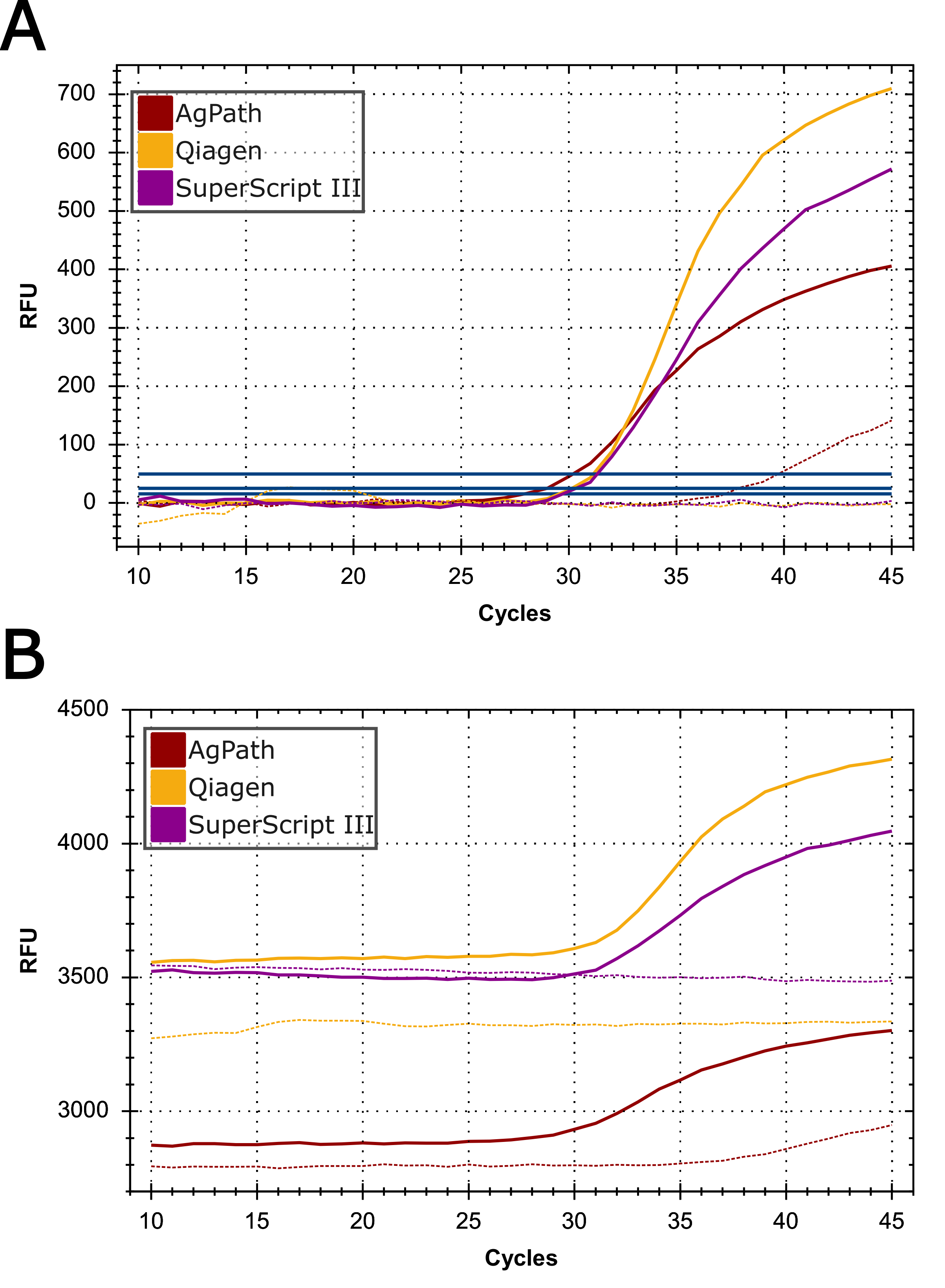

### Supplementary figure S04

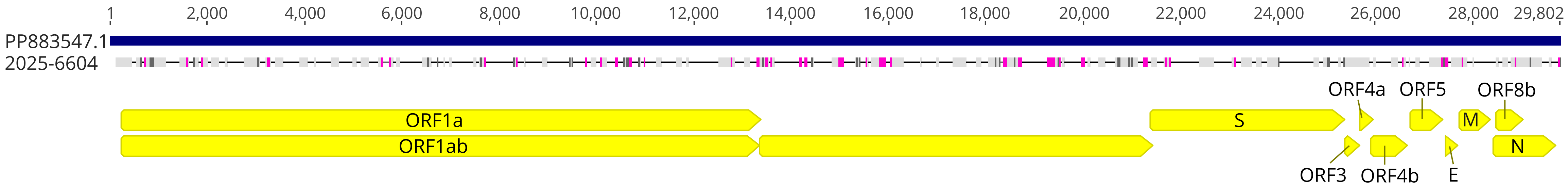

### Supplementary figure S05

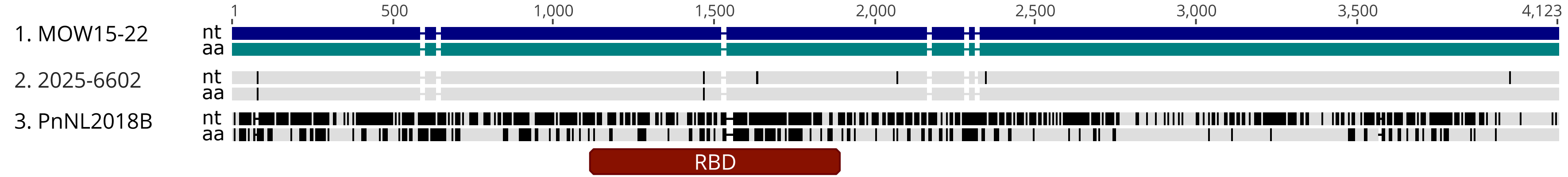
